# A novel ^18^F-labeled brain penetrant PET ligand for imaging poly(ADP-ribose) polymerase-1

**DOI:** 10.1101/2025.11.01.686021

**Authors:** Jimmy S. Patel, Xin Zhou, Jiahui Chen, Yabiao Gao, Chunyu Zhao, Zhendong Song, Taoqian Zhao, Qilong Hu, Xiaoyan Li, Chongjiao Li, Songyu Wu, David M. Schuster, Lily Yang, Yinlong Li, David S. Yu, Steven H. Liang

**Affiliations:** Department of Radiation Oncology, Winship Cancer Institute of Emory University, Atlanta, Georgia, 30322, United States; Department of Radiology and Imaging Sciences, Emory University, 1364 Clifton Road, Atlanta, Georgia 30322, United States; Department of Surgery, Winship Cancer Institute of Emory University, Atlanta, Georgia, 30322, United States; Wallace H. Coulter Department of Biomedical Engineering, Georgia Institute of Technology and Emory University, Atlanta, GA 30332, USA

**Keywords:** PARP, positron emission tomography, radioligand, breast, glioblastoma, prostate, pancreas

## Abstract

Poly(ADP-ribose) polymerase-1 (PARP-1) is a key mediator of DNA repair, and its inhibition has become a validated therapeutic strategy in homologous recombination–deficient cancers. However, tools for non-invasive assessment of PARP-1-specific expression remain limited. Here, we evaluated [^18^F]AZD9574, a next-generation PARP-1-selective PET radiotracer. [^18^F]AZD9574 was tested in a panel of breast, glioblastoma, prostate, and pancreatic cancer cell lines. Uptake correlated with PARP-1 expression and was dose-dependently blocked by a variety of clinically relevant PARP inhibitors, confirming its binding specificity. In 22Rv1 xenograft mouse models, the tracer demonstrated significant tumor accumulation that was specific to PARP-1, and *ex vivo* biodistribution confirmed organ uptake consistent with specific tumor binding and PARP-1 expression. Together, these findings establish [^18^F]AZD9574 as a promising PARP-1-targeted imaging agent with strong potential for advancing cancer research and therapeutic monitoring.

## Introduction

Poly(ADP-ribose) polymerase-1 (PARP-1) is a key DNA damage sensor that orchestrates repair of single-strand DNA breaks and has emerged as a validated therapeutic target in oncology.^1-3^ Tumors with homologous recombination deficient (HRD) pathways, such as BRCA-mutated cancers, demonstrate synthetic lethality with PARP inhibition, and multiple PARP inhibitors are now FDA-approved across ovarian, breast, and prostate cancers.^4-6^ Beyond oncology, PARP modulation has shown promise in inflammatory, cardiovascular, and neurologic disorders, underscoring its broad therapeutic relevance.^7-10^ These therapeutic effects are now understood to be driven primarily by PARP-1 modulation; however, current first-generation PARP inhibitors are not isoform-selective and inhibit multiple PARP family members.^11^ As a consequence, off-target PARP-2 inhibition has been linked to hematologic toxicity, including anemia.^12-15^ Despite their clinical success, major questions remain regarding PARP-1 expression patterns, real-time target engagement, and mechanisms of resistance *in vivo*, underscoring the need for noninvasive tools that can quantify these dimensions. Positron emission tomography (PET) imaging offers a powerful, non-invasive approach to study molecular targets such as PARP-1, yet existing tracers have significant limitations. Clinical-stage agents have demonstrated feasibility but are derived from PARP-1/2 non-selective inhibitors and show restricted blood–brain barrier (BBB) penetration, hindering their application in central nervous system (CNS) diseases and brain metastases.^16-21^ More recent ^11^C-labeled tracers demonstrate improved CNS permeability but face challenges of short half-life and unresolved issues of PARP subtype selectivity, limiting their translational potential.^22^

To overcome these barriers, we recently reported the development of [^18^F]AZD9574, a fluorine-18 isotopologue of the next-generation PARP-1-selective inhibitor AZD9574.^23^ In initial validation studies, [^18^F]AZD9574 demonstrated favorable pharmacokinetics, high specificity for PARP-1 over PARP-2, and the ability to cross the BBB, addressing key gaps that have limited prior tracers. Here, we expand upon these findings with comprehensive *in vitro* and *in vivo* evaluation across multiple oncologic model systems, including breast, prostate, pancreatic, and glioblastoma cell lines, as well as xenograft mouse models. These studies confirm the tracer’s selective accumulation in PARP-1-expressing tumors, demonstrate robust blockade with competitive inhibitors, and highlight favorable biodistribution and pharmacodynamics. Collectively, this work advances [^18^F]AZD9574 as a promising PARP-1-specific PET tracer with strong translational potential for studying PARP biology, monitoring therapeutic engagement, and guiding clinical decision-making in oncology.

## Methods

### Radiochemistry

The synthesis of [^18^F]AZD9574 followed the strategy we previously reported (**Figure S1**).^23^ Briefly, AZD9574, compound 5, contains a fluorine atom at a meta position of the picolinamide scaffold, enabling nucleophilic substitution without the need for additional activation groups. A brominated analog, compound 8, was designed as the radiolabeling precursor, and both compounds were synthesized in four steps from commercially available methyl picolinates. Compound 5 was obtained through a Pd-catalyzed Buchwald–Hartwig coupling, Boc deprotection, substitution with a quinoxalinone derivative, and amine–ester exchange. Compound 8 was prepared via analogous steps, with overall yields of 6% and 8%, respectively. As previously described, [^18^F]AZD9574 was synthesized from compound 8 via a direct SNAr displacement. Optimization of base, solvent, and temperature conditions identified DMSO at elevated temperature as most favorable, yielding [^18^F]AZD9574 with up to 25% radiochemical conversion. Under these conditions, the tracer was obtained in 15% non-decay corrected isolated yield, with excellent radiochemical purity (>99%) and high molar activity (>37 GBq/µmol).

### Western Blot

To extend our prior characterization of PARP-1 expression in PC3 prostate cancer cells, we performed western blot analysis across additional cancer cell lines. U87-MG (glioblastoma), 22Rv1 (prostate), PSN-1 (pancreatic), MDA-MB-436 (breast), and MDA-MB-231 (breast) cells were harvested and lysed in radioimmunoprecipitation assay (RIPA) lysis and extraction buffer (Thermo Scientific, 89901) supplemented with protease and phosphatase inhibitors (Thermo Scientific, A32965). Protein concentrations were quantified using the bicinchoninic acid (BCA) assay (Thermo Scientific, 23227). Equal amounts of protein were separated by SDS-PAGE, transferred to membranes, and probed with the following primary antibodies: PARP-1 polyclonal antibody (Cell Signaling Technology, 9532) and β-Tubulin monoclonal antibody (Invitrogen, MA5-16308-D680) as loading control. Detection was performed using Goat Anti-Rabbit IgG H&L (HRP) secondary antibody (Abcam, ab97080) and imaged on a ChemiDoc MP Imaging System (Bio-Rad). GAPDH (Abcam, ab181602) was used as the reference for U87-MG, 22RV1, and PSN-1 cell line while β-Tubulin was used as the reference loading control for the breast cancer cell lines.

### Cellular Uptake Studies

To extend our prior evaluation of [^18^F]AZD9574 uptake in PC3 cells, we conducted *in vitro* cellular uptake studies across the same panel of cancer cell lines as described in the Western blot. Cells were incubated with [^18^F]AZD9574 for up to 2 hours, and uptake was measured at defined timepoints. Blocking studies were performed using AZD9574, as well as clinically relevant PARP inhibitors (Olaparib, Rucaparib, Veliparib, Pamiparib), to confirm specificity. To assess selectivity for PARP-1 over PARP-2, the PARP-2-specific inhibitor UPF-1035 was also evaluated.

### Xenograft Production

Building on our prior experience with U87-MG and 22Rv1 xenograft models, J:nu outbred athymic nude mice (Jackson Laboratory) were used for xenograft production.^24^ Approximately 1 × 10^7 22Rv1 cells were resuspended in a 1:1 mixture of PBS and Matrigel and injected subcutaneously into the shoulder pad of each mouse to promote consistent tumor establishment. Tumor growth was monitored weekly by caliper measurements, and animals were enrolled for subsequent biodistribution and PET imaging studies with [^18^F]AZD9574 once tumors reached optimal size, typically 28–30 days post-implantation.

### Biodistribution

Based on prior studies, xenograft mice were sacrificed at 5, 15, 30, and 60 min after the administration of [^18^F]AZD9574 (20 µCi/100 µL) via the tail vein. Major organs of interest were collected, weighed, and measured by a gamma counter.^25, 26^

### Small Animal PET imaging

PET scans were acquired in tumor-bearing mice using a G8 scanner (Sofie). Animals were maintained under anesthesia with 1–2% isoflurane in air (v/v) throughout the imaging sessions. Approximately 40 µCi of [^18^F]AZD9574 was administered via intravenous injection through a preinstalled tail vein catheter. Static PET data were collected continuously for 60 minutes to 120 minutes. For blocking experiments, xenograft mice were pretreated with AZD9574 (1 mg/kg, i.v.) for 10 minutes prior to tracer administration. Image reconstruction and analysis were performed using AMIDE and PMOD software, as previously described, with regions and volumes of interest defined for quantitative assessment.^19, 27-30^

## Results

### Western Blot

Western blot analysis confirmed detectable PARP-1 expression in all tested cancer cell lines, with variable intensity across tumor types. Relative quantification revealed that human prostate carcinoma 22Rv1 and adenocarcinoma PSN-1 cells exhibited the highest PARP-1 expression compared to U87-MG glioma cells. Expression in MDA-MB-436 and MDA-MB-231 breast cancer cells was lower but still detectable when normalized to β-Tubulin. Additional analyses demonstrated that PARP-1 and PARP-2 were expressed at comparable levels, contrary to the conventional expectation of markedly lower PARP-2 expression and thereby highlighting inter-cell line heterogeneity (**Figure 1)**. These results extend our previous findings in prostatic adenocarcinoma PC3 cells by confirming robust PARP-1 expression in diverse tumor models, particularly in 22Rv1 and PSN-1 cells, which were subsequently prioritized for further radiotracer evaluation. Importantly, mapping PARP-1 expression across different tumor types not only informs the selection of preclinical models but also provides critical insight into which cancers may derive the greatest clinical benefit from PARP-1-targeted imaging and therapy.

**Figure 1:**
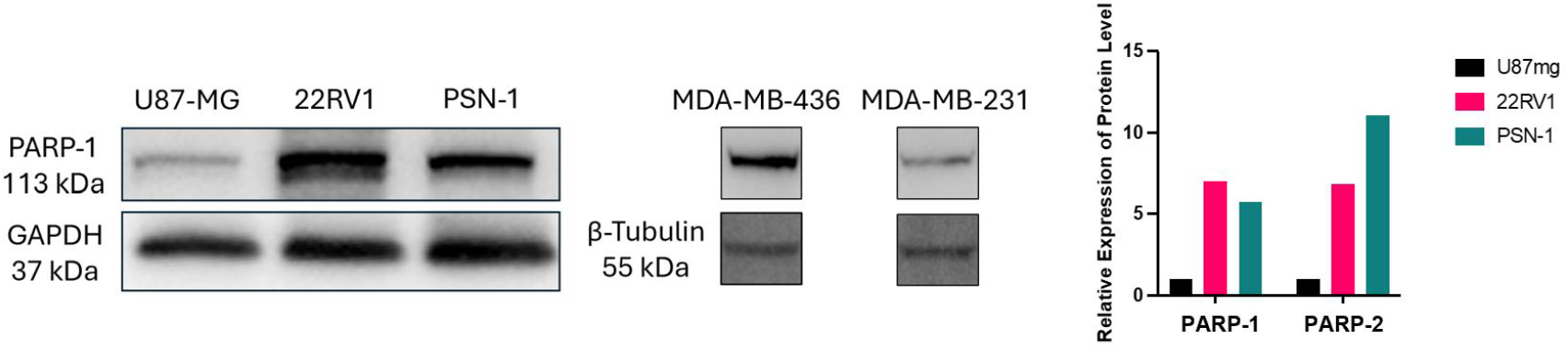
Western blot assessment of PARP-1 expression within U87-MG (glioblastoma), 22Rv1 (prostate), PSN-1 (pancreatic), MDA-MB-436 (breast), and MDA-MB-231 (breast) cancer cells. GAPDH was utilized as reference for U87-MG, 22RV1 and PSN-1; β-Tubulin was utilized as the standard for the latter two cell lines (MDA-MB-436 and MDA-MB-231). Right: Relative expression of PARP-1 and PARP-2 levels showing increased PARP-1 expression in both 22Rv1 and PSN-1 cells, relative to U87-MG cells.

### Cellular Uptake Studies

Radiotracer [^18^F]AZD9574 demonstrated time-dependent uptake across all tumor cell lines, with accumulation levels closely matching their PARP-1 expression profiles. Uptake was highest in 22Rv1 and PSN-1 cells, consistent with their strong PARP-1 expression, with relative uptake values at 2 hours of 45% and 1.4%, respectively, **Figure 2**. Blocking studies confirmed the tracer’s PARP-1 selectivity: AZD9574 and multiple PARP inhibitors significantly reduced uptake, while UPF-1035, a selective PARP-2 inhibitor, had no effect, **Figure 3**. Together, these findings reaffirm [^18^F]AZD9574 as a PARP-1-specific tracer with high binding affinity and selectivity *in vitro*. Based on these results, 22Rv1 cells were prioritized for subsequent tumor xenograft studies.

**Figure 2:**
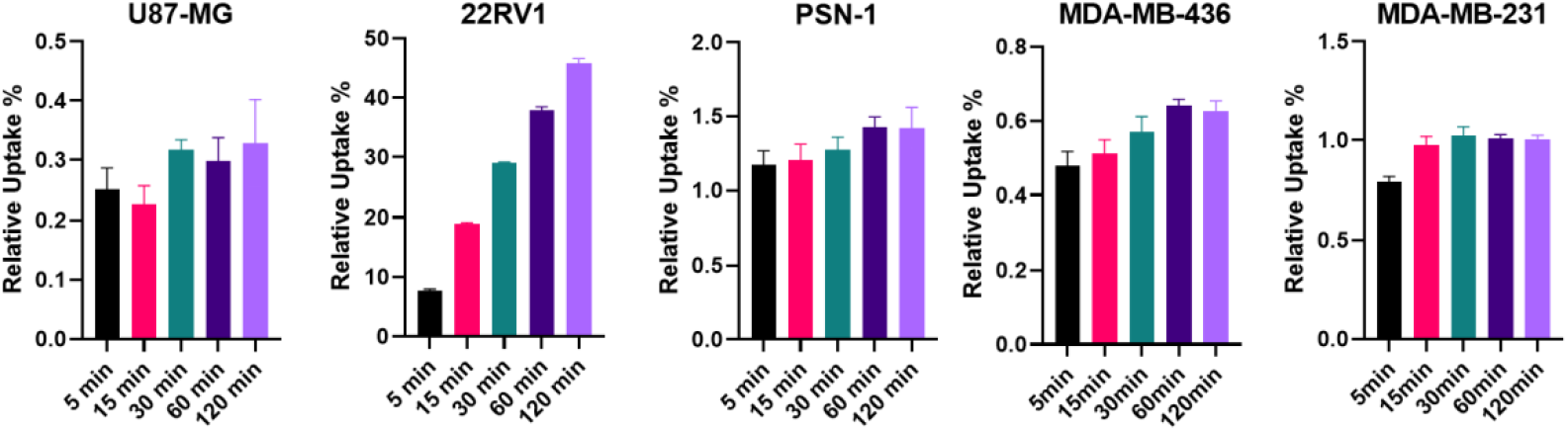
Uptake of [^18^F]AZD9574 within the aforementioned cell lines. Intracellular concentrations of [^18^F]AZD9574 was generally found to increase in a time-dependent manner across 120 minutes, with the highest intracellular concentrations found within 22Rv1 cells and PSN-1 cells.

**Figure 3:**
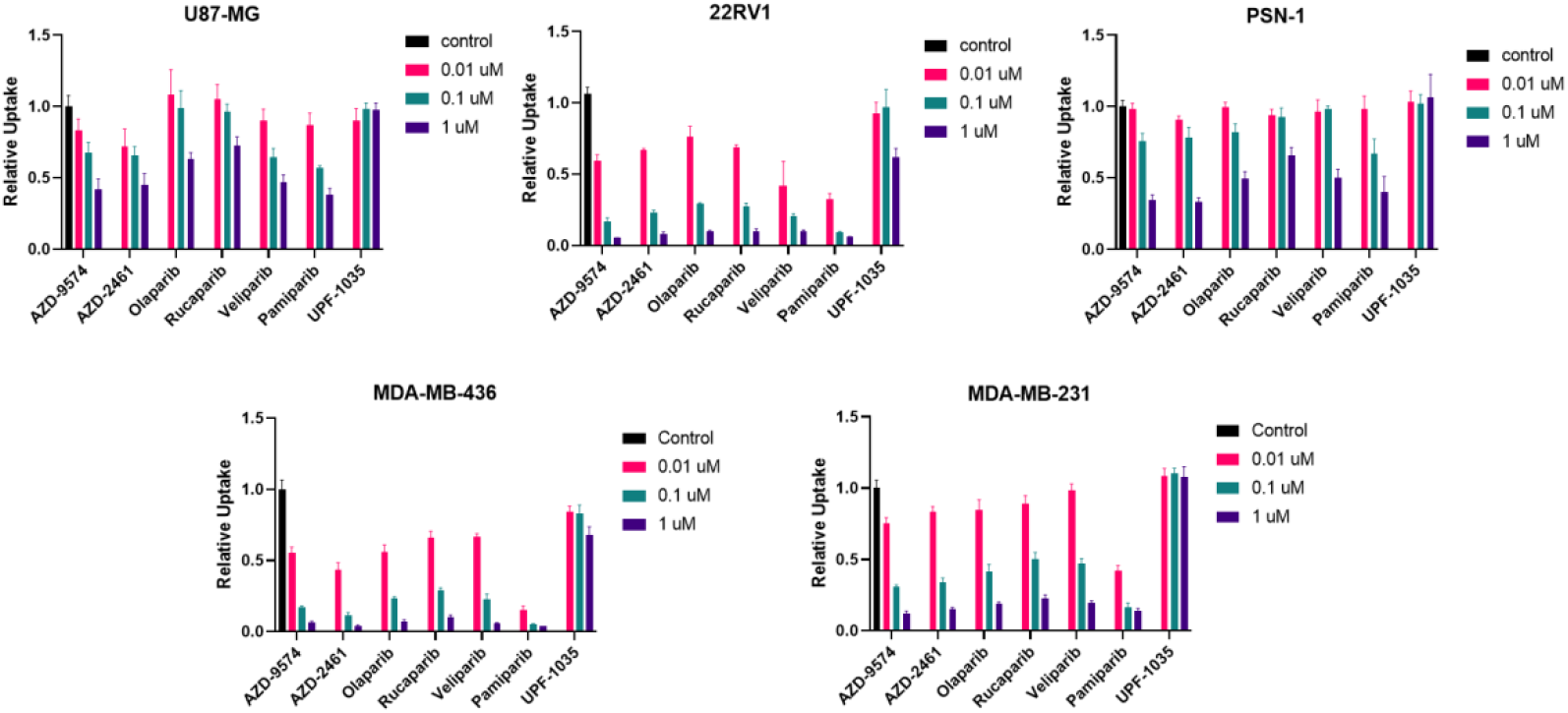
Intracellular uptake of [^18^F]AZD9574 was effectively reduced in a dose-dependent manner upon treatment with blocking agents. Across all tested cell lines, blocking with 0.01 µM, 0.1 µM, and 1 µM of a PARP inhibitor decreased radioligand uptake, indicating high specific binding *in vitro*. In contrast, UPF-1035, a PARP-2 specific inhibitor, failed to reduce cellular uptake, further confirming the PARP-1 selectivity of [^18^F]AZD9574.

### Biodistribution

*Ex vivo* biodistribution studies demonstrated rapid and widespread distribution of [^18^F]AZD9574 following intravenous administration. Radioactivity was detected in multiple organs, with patterns consistent with previously reported pharmacokinetic profiles of AZD9574.^23, 31^ Notably, significant tracer accumulation was observed in tumor tissue, supporting tumor-specific uptake. Tracer levels in blood decreased progressively over time, while clearance through hepatobiliary and renal pathways was evident from measurable activity in the liver, kidneys, and intestines. The high uptake seen within the spleen, pancreas, and other organs of the gastrointestinal tract are consistent with prior reports studying PARP radiotracer *in vivo* **Figure 4**.^19, 32^

**Figure 4:**
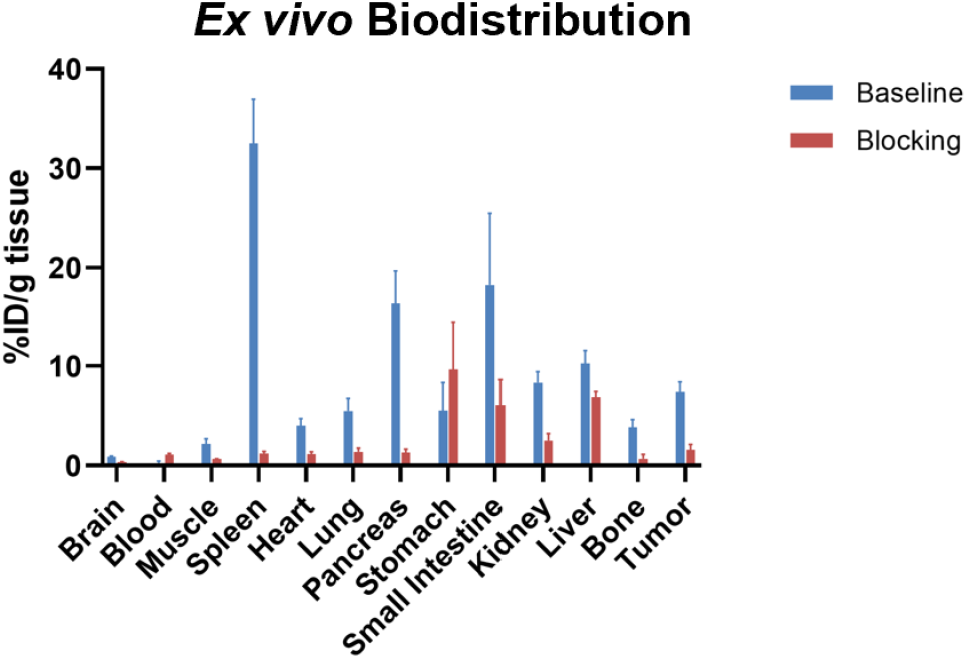
*Ex vivo* biodistribution analysis in 22Rv1 tumor mice (60 min post injection, n = 5 for both baseline and blocking study). For the blocking study, AZD9574 was introduced as the blocking agent 10 mins prior to radiotracer injection.

Pre-treatment with unlabeled AZD9574 markedly reduced tumor uptake, confirming competitive inhibition at the PARP-1 binding site. This blocking effect was tumor-selective, with non-target organs showing relatively unchanged distribution profiles, further supporting the specificity of [^18^F]AZD9574 for PARP-1. Collectively, these biodistribution data validate both the *in vivo* stability of the tracer and its ability to selectively accumulate in PARP-1-expressing tumors.

### Small Animal PET Imaging

Small animal PET imaging of mice bearing 22Rv1 xenografts demonstrated clear *in vivo* tracer accumulation with favorable pharmacokinetics. The maximum standardized uptake value (SUV) reached 0.96 at 1-hour post-injection and decreased to 0.76 at 2 hours (n = 3), consistent with stable tumor retention and gradual clearance over time, **Figure 5**. As expected, pre-administration of unlabeled AZD9574 appreciably reduced tumor uptake, with a marked decrease in maximum SUV (n = 4), confirming the specificity of [^18^F]AZD9574 binding *in vivo*.

**Figure 5:**
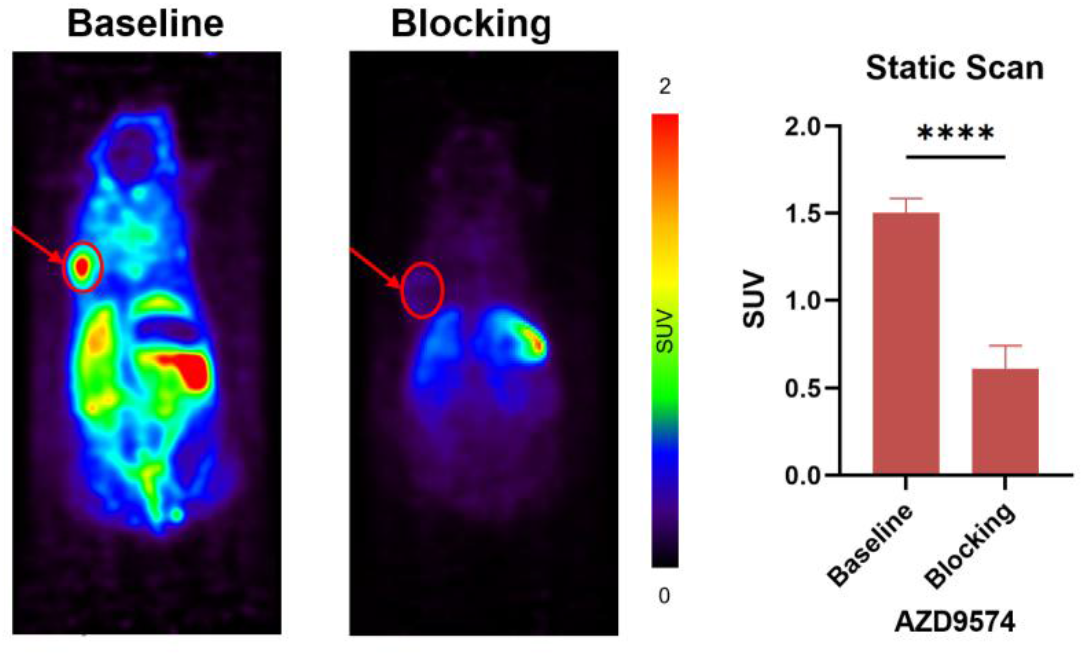
Small animal PET imaging of 22Rv1 xenograft mice (left panel). Blocking with AZD9574 appreciably decreases maximum SUV in 22Rv1 xenograft mice (n = 4, right panel). All data are mean ± SD when applicable. **** = p ≤ 0.0001

## Discussion

PARP inhibitors (PARPi) have emerged as a major therapeutic option for patients with HRD cancers, with demonstrated benefit in ovarian, breast, prostate, and pancreatic malignancies.^4-6^ Despite these advances, clinical responses are heterogeneous. Even among patients selected for BRCA1/2 or other HRD gene mutations, only a fraction respond, while others with no identifiable HRD still derive benefit.^33, 34^ This heterogeneity highlights the need for biomarkers that extend beyond genetic profiling and provide direct, dynamic measures of target expression and drug engagement. PARP-1 is the predominant isoform implicated in the synthetic lethal interaction with BRCA deficiency, yet it has never been systematically evaluated *in vivo* across patients. Noninvasive imaging approaches, particularly PET with radiolabeled PARPi analogs, provide a powerful means of quantifying PARP-1 expression and distribution across tumors and metastatic sites, potentially guiding therapy decisions where conventional assays fall short.^19, 35-37^

Several early studies have demonstrated the feasibility of PARP-targeted PET imaging. [^18^F]FluorThanatrace has advanced into first-in-human imaging and shown correlation with PARP-1 expression, response to PARPi therapy, and competitive blocking by therapeutic PARPi agents. These findings underscore its potential as a predictive biomarker and a tool for monitoring target engagement.^16, 37^ More broadly, PET imaging offers advantages uniquely suited to precision oncology which includes the ability to provide whole-body disease mapping, to capture intra-patient/tumoral heterogeneity, and to quantify pharmacokinetics and pharmacodynamics of targeted therapies. This concept parallels the success of tracers such as [^18^F]fluoroestradiol for breast cancer, PSMA tracers for prostate cancer, which are now FDA-approved and routinely applied in clinical practice.^38-40^ Despite the promise, limitations of existing PARP inhibitor based tracers remain significant. Most available agents, are derived from non-selective PARP-1/2 inhibitors, raising concerns about off-target binding and interpretability of imaging signals. In addition, their limited blood-brain barrier penetration restricts application in CNS metastasis and primary brain lesions, such as gliomas, which represent important and growing indications for PARPi therapy.^41^

The development of [^18^F]AZD9574 directly addresses several of these unmet needs. As an isotopologue of AZD9574, a next-generation PARP-1-selective inhibitor currently in clinical trials (NCT05417594), [^18^F]AZD9574 offers high selectivity for PARP-1 over PARP-2 and demonstrated ability to penetrate the blood-brain barrier.^23, 42, 43^ Our findings show that the tracer exhibits time-dependent, PARP-1-specific uptake across multiple tumor models, with competitive blocking studies confirming specificity.^44^ *Ex vivo* biodistribution demonstrated notable tumor uptake consistent with PARP-1 expression, while *in vivo* PET imaging in tumor xenografts confirmed favorable pharmacokinetics and reduced uptake upon blockade. Together, these data support the advancement of [^18^F]AZD9574 as a translational radiotracer with broad potential applications. While the present study primarily establishes tumor selectivity and *in vivo* pharmacokinetics, our prior work demonstrating BBB penetration of [^18^F]AZD9574 suggests additional promise for applications in CNS malignancies where earlier tracers have struggled.^23^

Another critical dimension to the utility of PARP imaging lies in the context of resistance. While PARPi are effective in many HRD cancers, resistance is nearly universal with prolonged therapy. Several mechanisms have been elucidated, including restoration of homologous recombination via secondary reversion mutations in BRCA1/2 or other HR genes, stabilization of replication forks to avoid collapse, mutations in PARP-1 itself that alter drug binding or trapping, and upregulation of efflux pumps such as P-glycoprotein that reduce intracellular drug exposure. Epigenetic reprogramming and altered expression of DNA damage response proteins further complicate the landscape.^33, 34, 45-47^ These resistance pathways mean that HRD status alone cannot reliably predict long-term benefit from PARPi. In this setting, noninvasive imaging of PARP-1 has unique potential to provide real-time insights into whether tumors remain targetable during therapy. For instance, declining tracer uptake could reflect reduced PARP-1 expression or accessibility, signaling resistance, while persistent uptake may identify lesions that remain vulnerable to inhibition. By coupling imaging with genomic assays, resistance mechanisms could be stratified *in vivo*, offering opportunities to tailor therapy adaptively.

Despite the promising data presented here, important limitations remain. As with all small animal models, xenografts cannot fully replicate the complexity of human tumors, including the microenvironment and immune interactions. While our current and previous study has shown serum stability within *in vitro* and *in vivo* small rodent systems, metabolite formation and tracer stability will need to be rigorously assessed in nonhuman primates and early human studies to confirm that the imaging signal reflects intact [^18^F]AZD9574. Additionally, cancers with low baseline PARP-1 expression may yield low tumor-to-background ratios, potentially limiting sensitivity in specific contexts. Finally, standardization of imaging protocols, quantification methods, and correlation with tissue-based assays will be essential for clinical translation.

## Conclusion

Looking forward, [^18^F]AZD9574 holds significant translational potential. Its high PARP-1 selectivity and BBB penetration position it uniquely among existing PARP tracers. Early-phase clinical studies will be required to establish safety, dosimetry, pharmacokinetics, and correlation of uptake with PARP-1 expression. Beyond baseline imaging, longitudinal studies could assess target engagement and emergence of resistance during therapy. Ultimately, PARP-1 imaging could be integrated into theranostic frameworks, guiding patient selection, optimizing drug dosing, and improving outcomes by enabling truly personalized use of PARPi.

## Acknowledgments

We thank Emory Center for Systems Imaging Radiopharmacy (Ronald J. Crowe, RPh, BCNP; Karen Dolph, RPh; M. Shane Waldrep) & Department of Radiology and Imaging Sciences, Emory University School of Medicine for general support. We also thank the National Institute of Mental Health’s Psychoactive Drug Screening Program (NIMH PDSP) for compound off-target screening. J.S.P. is supported by NCI T32CA275777 and the recipient of 2025 Elkin Fellowship. S.H.L. gratefully acknowledges the support provided, in part, by the NIH grant (AG094161), Emory Radiology Chair Fund, and Emory School of Medicine Endowed Directorship. We thank Emory Center for Systems Imaging for assistance with small animal micro-PET/CT system, which was funded by the NIH grant (S10OD034326) and Emory Integrated Core Facilities.

## References

1. Ray Chaudhuri A, Nussenzweig A. The multifaceted roles of PARP-1 in DNA repair and chromatin remodelling. Nat Rev Mol Cell Biol. 2017;18(10):610–21. Epub 20170705. doi: 10.1038/nrm.2017.53. PubMed PMID: 28676700; PMCID: PMC6591728.

2. Ko HL, Ren EC. Functional Aspects of PARP-1 in DNA Repair and Transcription. Biomolecules. 2012;2(4):524–48. Epub 20121112. doi: 10.3390/biom2040524. PubMed PMID: 24970148; PMCID: PMC4030864.

3. Luo X, Kraus WL. On PAR with PARP: cellular stress signaling through poly(ADP-ribose) and PARP-1. Genes Dev. 2012;26(5):417–32. doi: 10.1101/gad.183509.111. PubMed PMID: 22391446; PMCID: PMC3305980.

4. Fizazi K, Piulats JM, Reaume MN, Ostler P, McDermott R, Gingerich JR, Pintus E, Sridhar SS, Bambury RM, Emmenegger U, Lindberg H, Morris D, Nole F, Staffurth J, Redfern C, Saez MI, Abida W, Daugaard G, Heidenreich A, Krieger L, Sautois B, Loehr A, Despain D, Heyes CA, Watkins SP, Chowdhury S, Ryan CJ, Bryce AH, Investigators T. Rucaparib or Physician’s Choice in Metastatic Prostate Cancer. N Engl J Med. 2023;388(8):719–32. Epub 20230216. doi: 10.1056/NEJMoa2214676. PubMed PMID: 36795891; PMCID: PMC10064172.

5. Gonzalez-Martin A, Pothuri B, Vergote I, DePont Christensen R, Graybill W, Mirza MR, McCormick C, Lorusso D, Hoskins P, Freyer G, Baumann K, Jardon K, Redondo A, Moore RG, Vulsteke C, O’Cearbhaill RE, Lund B, Backes F, Barretina-Ginesta P, Haggerty AF, Rubio-Perez MJ, Shahin MS, Mangili G, Bradley WH, Bruchim I, Sun K, Malinowska IA, Li Y, Gupta D, Monk BJ, Investigators PE-OG-. Niraparib in Patients with Newly Diagnosed Advanced Ovarian Cancer. N Engl J Med. 2019;381(25):2391–402. Epub 20190928. doi: 10.1056/NEJMoa1910962. PubMed PMID: 31562799.

6. Robson M, Im SA, Senkus E, Xu B, Domchek SM, Masuda N, Delaloge S, Li W, Tung N, Armstrong A, Wu W, Goessl C, Runswick S, Conte P. Olaparib for Metastatic Breast Cancer in Patients with a Germline BRCA Mutation. N Engl J Med. 2017;377(6):523–33. Epub 20170604. doi: 10.1056/NEJMoa1706450. PubMed PMID: 28578601.

7. Curtin NJ, Szabo C. Poly(ADP-ribose) polymerase inhibition: past, present and future. Nat Rev Drug Discov. 2020;19(10):711–36. Epub 20200903. doi: 10.1038/s41573-020-0076-6. PubMed PMID: 32884152.

8. Kim C, Chen C, Yu Y. Avoid the trap: Targeting PARP-1 beyond human malignancy. Cell Chem Biol. 2021;28(4):456–62. Epub 20210302. doi: 10.1016/j.chembiol.2021.02.004. PubMed PMID: 33657415; PMCID: PMC8052287.

9. Pazzaglia S, Pioli C. Multifaceted Role of PARP-1 in DNA Repair and Inflammation: Pathological and Therapeutic Implications in Cancer and Non-Cancer Diseases. Cells. 2019;9(1). Epub 20191222. doi: 10.3390/cells9010041. PubMed PMID: 31877876; PMCID: PMC7017201.

10. Thapa K, Khan H, Sharma U, Grewal AK, Singh TG. Poly (ADP-ribose) polymerase-1 as a promising drug target for neurodegenerative diseases. Life Sci. 2021;267:118975. Epub 20201231. doi: 10.1016/j.lfs.2020.118975. PubMed PMID: 33387580.

11. Wahlberg E, Karlberg T, Kouznetsova E, Markova N, Macchiarulo A, Thorsell AG, Pol E, Frostell A, Ekblad T, Oncu D, Kull B, Robertson GM, Pellicciari R, Schuler H, Weigelt J. Family-wide chemical profiling and structural analysis of PARP and tankyrase inhibitors. Nat Biotechnol. 2012;30(3):283–8. Epub 20120219. doi: 10.1038/nbt.2121. PubMed PMID: 22343925.

12. LaFargue CJ, Dal Molin GZ, Sood AK, Coleman RL. Exploring and comparing adverse events between PARP inhibitors. Lancet Oncol. 2019;20(1):e15–e28. doi: 10.1016/S1470-2045(18)30786-1. PubMed PMID: 30614472; PMCID: PMC7292736.

13. Coleman RL, Oza AM, Lorusso D, Aghajanian C, Oaknin A, Dean A, Colombo N, Weberpals JI, Clamp A, Scambia G, Leary A, Holloway RW, Gancedo MA, Fong PC, Goh JC, O’Malley DM, Armstrong DK, Garcia-Donas J, Swisher EM, Floquet A, Konecny GE, McNeish IA, Scott CL, Cameron T, Maloney L, Isaacson J, Goble S, Grace C, Harding TC, Raponi M, Sun J, Lin KK, Giordano H, Ledermann JA, investigators A. Rucaparib maintenance treatment for recurrent ovarian carcinoma after response to platinum therapy (ARIEL3): a randomised, double-blind, placebo-controlled, phase 3 trial. Lancet. 2017;390(10106):1949–61. Epub 20170912. doi: 10.1016/S0140-6736(17)32440-6. PubMed PMID: 28916367; PMCID: PMC5901715.

14. Mirza MR, Monk BJ, Herrstedt J, Oza AM, Mahner S, Redondo A, Fabbro M, Ledermann JA, Lorusso D, Vergote I, Ben-Baruch NE, Marth C, Madry R, Christensen RD, Berek JS, Dorum A, Tinker AV, du Bois A, Gonzalez-Martin A, Follana P, Benigno B, Rosenberg P, Gilbert L, Rimel BJ, Buscema J, Balser JP, Agarwal S, Matulonis UA, Investigators E-ON. Niraparib Maintenance Therapy in Platinum-Sensitive, Recurrent Ovarian Cancer. N Engl J Med. 2016;375(22):2154–64. Epub 20161007. doi: 10.1056/NEJMoa1611310. PubMed PMID: 27717299.

15. Pujade-Lauraine E, Ledermann JA, Selle F, Gebski V, Penson RT, Oza AM, Korach J, Huzarski T, Poveda A, Pignata S, Friedlander M, Colombo N, Harter P, Fujiwara K, Ray-Coquard I, Banerjee S, Liu J, Lowe ES, Bloomfield R, Pautier P, investigators SOE-O. Olaparib tablets as maintenance therapy in patients with platinum-sensitive, relapsed ovarian cancer and a BRCA1/2 mutation (SOLO2/ENGOT-Ov21): a double-blind, randomised, placebo-controlled, phase 3 trial. Lancet Oncol. 2017;18(9):1274–84. Epub 20170725. doi: 10.1016/S1470-2045(17)30469-2. PubMed PMID: 28754483.

16. Lee HS, Schwarz SW, Schubert EK, Chen DL, Doot RK, Makvandi M, Lin LL, McDonald ES, Mankoff DA, Mach RH. The Development of (18)F Fluorthanatrace: A PET Radiotracer for Imaging Poly (ADP-Ribose) Polymerase-1. Radiol Imaging Cancer. 2022;4(1):e210070. doi: 10.1148/rycan.210070. PubMed PMID: 35089089; PMCID: PMC8830434.

17. Schoder H, Franca PDS, Nakajima R, Burnazi E, Roberts S, Brand C, Grkovski M, Mauguen A, Dunphy MP, Ghossein RA, Lyashchenko SK, Lewis JS, O’Donoghue JA, Ganly I, Patel SG, Lee NY, Reiner T. Safety and Feasibility of PARP-1/2 Imaging with (18)F-PARPi in Patients with Head and Neck Cancer. Clin Cancer Res. 2020;26(13):3110–6. Epub 20200403. doi: 10.1158/1078-0432.CCR-19-3484. PubMed PMID: 32245901; PMCID: PMC7421489.

18. Tong J, Chen B, Tan PW, Kurpiewski S, Cai Z. Poly (ADP-ribose) polymerases as PET imaging targets for central nervous system diseases. Front Med (Lausanne). 2022;9:1062432. Epub 20221110. doi: 10.3389/fmed.2022.1062432. PubMed PMID: 36438061; PMCID: PMC9685622.

19. Wilson TC, Xavier MA, Knight J, Verhoog S, Torres JB, Mosley M, Hopkins SL, Wallington S, Allen PD, Kersemans V, Hueting R, Smart S, Gouverneur V, Cornelissen B. PET Imaging of PARP Expression Using (18)F-Olaparib. J Nucl Med. 2019;60(4):504–10. Epub 20181102. doi: 10.2967/jnumed.118.213223. PubMed PMID: 30389822; PMCID: PMC6448459.

20. Deng X, Rong J, Wang L, Vasdev N, Zhang L, Josephson L, Liang SH. Chemistry for Positron Emission Tomography: Recent Advances in (11) C-, (18) F-, (13) N-, and (15) O-Labeling Reactions. Angew Chem Int Ed Engl. 2019;58(9):2580–605. Epub 20190114. doi: 10.1002/anie.201805501. PubMed PMID: 30054961; PMCID: PMC6405341.

21. Rong J, Haider A, Jeppesen TE, Josephson L, Liang SH. Radiochemistry for positron emission tomography. Nat Commun. 2023;14(1):3257. Epub 20230605. doi: 10.1038/s41467-023-36377-4. PubMed PMID: 37277339; PMCID: PMC10241151.

22. Chen B, Ojha DP, Toyonaga T, Tong J, Pracitto R, Thomas MA, Liu M, Kapinos M, Zhang L, Zheng MQ, Holden D, Fowles K, Ropchan J, Nabulsi N, De Feyter H, Carson RE, Huang Y, Cai Z. Preclinical evaluation of a brain penetrant PARP PET imaging probe in rat glioblastoma and nonhuman primates. Eur J Nucl Med Mol Imaging. 2023;50(7):2081–99. Epub 20230228. doi: 10.1007/s00259-023-06162-y. PubMed PMID: 36849748.

23. Zhou X, Chen J, Patel JS, Ran W, Li Y, Van RS, Ibrahim MMH, Zhao C, Gao Y, Rong J, Chaudhary AF, Li G, Hu J, Davenport AT, Daunais JB, Shao Y, Ran C, Collier TL, Haider A, Schuster DM, Levey AI, Wang L, Corfas G, Liang SH. Imaging poly(ADP-ribose) polymerase-1 (PARP-1) in vivo with 18F-labeled brain penetrant positron emission tomography (PET) ligand. Acta Pharmaceutica Sinica B. 2025. doi: 10.1016/j.apsb.2025.05.020.

24. Zhao C, Rong J, Crowe R, Collier L, Liang S. Development of a novel 18F-labeled PET ligand for imaging fibroblast activation protein. Journal of Nuclear Medicine. 2024;65(Supplement 2):241810-.

25. Rong J, Zhao C, Chaudhary AF, Chen J, Zhou X, Zhang K, Song Z, Sun Z, Gao Y, Zhang Z, Feng S, Collier TL, Yuan H, Patel JS, Haider A, Li Y, Liang SH. Development of a Novel (18)F-Labeled Radioligand for Imaging Cholesterol 24-Hydroxylase with Positron Emission Tomography. ACS Pharmacol Transl Sci. 2025;8(3):800–7. Epub 20250224. doi: 10.1021/acsptsci.4c00683. PubMed PMID: 40109739; PMCID: PMC11915032.

26. Rong J, Zhao C, Chaudhary AF, Jones E, Van R, Song Z, Li Y, Chen J, Zhou X, Patel JS, Gao Y, Sun Z, Feng S, Zhang Z, Collier TL, Ran C, Haider A, Shao Y, Yuan H, Liang SH. Development of a Novel (18)F-Labeled Radioligand for Imaging Phosphodiesterase 7 with Positron Emission Tomography. Mol Pharm. 2025;22(3):1657–66. Epub 20250219. doi: 10.1021/acs.molpharmaceut.4c01379. PubMed PMID: 39970438; PMCID: PMC11881136.

27. Chen Z, Mori W, Deng X, Cheng R, Ogasawara D, Zhang G, Schafroth MA, Dahl K, Fu H, Hatori A, Shao T, Zhang Y, Yamasaki T, Zhang X, Rong J, Yu Q, Hu K, Fujinaga M, Xie L, Kumata K, Gou Y, Chen J, Gu S, Bao L, Wang L, Collier TL, Vasdev N, Shao Y, Ma JA, Cravatt BF, Fowler C, Josephson L, Zhang MR, Liang SH. Design, Synthesis, and Evaluation of Reversible and Irreversible Monoacylglycerol Lipase Positron Emission Tomography (PET) Tracers Using a “Tail Switching” Strategy on a Piperazinyl Azetidine Skeleton. J Med Chem. 2019;62(7):3336–53. Epub 20190321. doi: 10.1021/acs.jmedchem.8b01778. PubMed PMID: 30829483; PMCID: PMC6581563.

28. Chen Z, Mori W, Zhang X, Yamasaki T, Dunn PJ, Zhang G, Fu H, Shao T, Zhang Y, Hatori A, Ma L, Fujinaga M, Xie L, Deng X, Li H, Yu Q, Rong J, Josephson L, Ma JA, Shao Y, Tomita S, Zhang MR, Liang SH. Synthesis, pharmacology and preclinical evaluation of (11)C-labeled 1,3-dihydro-2H-benzo[d]imidazole-2-ones for imaging gamma8-dependent transmembrane AMPA receptor regulatory protein. Eur J Med Chem. 2018;157:898–908. Epub 20180809. doi: 10.1016/j.ejmech.2018.08.019. PubMed PMID: 30145376; PMCID: PMC6245653.

29. Cheng R, Mori W, Ma L, Alhouayek M, Hatori A, Zhang Y, Ogasawara D, Yuan G, Chen Z, Zhang X, Shi H, Yamasaki T, Xie L, Kumata K, Fujinaga M, Nagai Y, Minamimoto T, Svensson M, Wang L, Du Y, Ondrechen MJ, Vasdev N, Cravatt BF, Fowler C, Zhang MR, Liang SH. In Vitro and in Vivo Evaluation of (11)C-Labeled Azetidinecarboxylates for Imaging Monoacylglycerol Lipase by PET Imaging Studies. J Med Chem. 2018;61(6):2278–91. Epub 20180309. doi: 10.1021/acs.jmedchem.7b01400. PubMed PMID: 29481079; PMCID: PMC5966020.

30. Liang SH, Southon AG, Fraser BH, Krause-Heuer AM, Zhang B, Shoup TM, Lewis R, Volitakis I, Han Y, Greguric I, Bush AI, Vasdev N. Novel Fluorinated 8-Hydroxyquinoline Based Metal Ionophores for Exploring the Metal Hypothesis of Alzheimer’s Disease. ACS Med Chem Lett. 2015;6(9):1025–9. Epub 20150810. doi: 10.1021/acsmedchemlett.5b00281. PubMed PMID: 26396692; PMCID: PMC4569883.

31. Zhou X CJ, Patel JS, Ran W, Li Y, Van RS, et al. Imaging poly(ADP-ribose) polymerase-1 (PARP-1) in vivo with 18F-labeled brain penetrant positron emission tomography (PET) ligand.2024. doi: 10.26434/chemrxiv-2024-8pqmg-v2.

32. Michel LS, Dyroff S, Brooks FJ, Spayd KJ, Lim S, Engle JT, Phillips S, Tan B, Wang-Gillam A, Bognar C, Chu W, Zhou D, Mach RH, Laforest R, Chen DL. PET of Poly (ADP-Ribose) Polymerase Activity in Cancer: Preclinical Assessment and First In-Human Studies. Radiology. 2017;282(2):453–63. Epub 20161114. doi: 10.1148/radiol.2016161929. PubMed PMID: 27841728; PMCID: PMC5283874.

33. Dias MP, Moser SC, Ganesan S, Jonkers J. Understanding and overcoming resistance to PARP inhibitors in cancer therapy. Nat Rev Clin Oncol. 2021;18(12):773–91. Epub 20210720. doi: 10.1038/s41571-021-00532-x. PubMed PMID: 34285417.

34. Li H, Liu ZY, Wu N, Chen YC, Cheng Q, Wang J. PARP inhibitor resistance: the underlying mechanisms and clinical implications. Mol Cancer. 2020;19(1):107. Epub 20200620. doi: 10.1186/s12943-020-01227-0. PubMed PMID: 32563252; PMCID: PMC7305609.

35. Makvandi M, Pantel A, Schwartz L, Schubert E, Xu K, Hsieh CJ, Hou C, Kim H, Weng CC, Winters H, Doot R, Farwell MD, Pryma DA, Greenberg RA, Mankoff DA, Simpkins F, Mach RH, Lin LL. A PET imaging agent for evaluating PARP-1 expression in ovarian cancer. J Clin Invest. 2018;128(5):2116–26. Epub 20180416. doi: 10.1172/JCI97992. PubMed PMID: 29509546; PMCID: PMC5919879.

36. Makvandi M, Xu K, Lieberman BP, Anderson RC, Effron SS, Winters HD, Zeng C, McDonald ES, Pryma DA, Greenberg RA, Mach RH. A Radiotracer Strategy to Quantify PARP-1 Expression In Vivo Provides a Biomarker That Can Enable Patient Selection for PARP Inhibitor Therapy. Cancer Res. 2016;76(15):4516–24. Epub 20160603. doi: 10.1158/0008-5472.CAN-16-0416. PubMed PMID: 27261505; PMCID: PMC5549277.

37. Pantel AR, Gitto SB, Makvandi M, Kim H, Medvedv S, Weeks JK, Torigian DA, Hsieh CJ, Ferman B, Latif NA, Tanyi JL, Martin LP, Lanzo SM, Liu F, Cao Q, Mills GB, Doot RK, Mankoff DA, Mach RH, Lin LL, Simpkins F. [18F]FluorThanatrace ([18F]FTT) PET Imaging of PARP-Inhibitor Drug-Target Engagement as a Biomarker of Response in Ovarian Cancer, a Pilot Study. Clin Cancer Res. 2023;29(8):1515–27. doi: 10.1158/1078-0432.CCR-22-1602. PubMed PMID: 36441795; PMCID: PMC12012890.

38. Giesel FL, Knorr K, Spohn F, Will L, Maurer T, Flechsig P, Neels O, Schiller K, Amaral H, Weber WA, Haberkorn U, Schwaiger M, Kratochwil C, Choyke P, Kramer V, Kopka K, Eiber M. Detection Efficacy of (18)F-PSMA-1007 PET/CT in 251 Patients with Biochemical Recurrence of Prostate Cancer After Radical Prostatectomy. J Nucl Med. 2019;60(3):362–8. Epub 20180724. doi: 10.2967/jnumed.118.212233. PubMed PMID: 30042163; PMCID: PMC6424235.

39. O’Brien SR, Edmonds CE, Lanzo SM, Weeks JK, Mankoff DA, Pantel AR. (18)F-Fluoroestradiol: Current Applications and Future Directions. Radiographics. 2023;43(3):e220143. doi: 10.1148/rg.220143. PubMed PMID: 36821506.

40. Parent EE, Schuster DM. Update on (18)F-Fluciclovine PET for Prostate Cancer Imaging. J Nucl Med. 2018;59(5):733–9. Epub 20180309. doi: 10.2967/jnumed.117.204032. PubMed PMID: 29523631; PMCID: PMC6910635.

41. Sim HW, Galanis E, Khasraw M. PARP Inhibitors in Glioma: A Review of Therapeutic Opportunities. Cancers (Basel). 2022;14(4). Epub 20220216. doi: 10.3390/cancers14041003. PubMed PMID: 35205750; PMCID: PMC8869934.

42. Johannes JW, Balazs AYS, Barratt D, Bista M, Chuba MD, Cosulich S, Critchlow SE, Degorce SL, Di Fruscia P, Edmondson SD, Embrey KJ, Fawell S, Ghosh A, Gill SJ, Gunnarsson A, Hande SM, Heightman TD, Hemsley P, Illuzzi G, Lane J, Larner CJB, Leo E, Liu L, Madin A, McWilliams L, O’Connor MJ, Orme JP, Pachl F, Packer MJ, Pei X, Pike A, Schimpl M, She H, Staniszewska AD, Talbot V, Underwood E, Varnes JG, Xue L, Yao T, Zhang K, Zhang AX, Zheng X. Discovery of 6-Fluoro-5-4-[(5-fluoro-2-methyl-3-oxo-3,4-dihydroquinoxalin-6-yl)methyl]piperazin-1-yl-N-methylpyridine-2-carboxamide (AZD9574): A CNS-Penetrant, PARP-1-Selective Inhibitor. J Med Chem. 2024;67(24):21717–28. Epub 20241210. doi: 10.1021/acs.jmedchem.4c01725. PubMed PMID: 39655996.

43. Staniszewska AD, Pilger D, Gill SJ, Jamal K, Bohin N, Guzzetti S, Gordon J, Hamm G, Mundin G, Illuzzi G, Pike A, McWilliams L, Maglennon G, Rose J, Hawthorne G, Cortes Gonzalez M, Halldin C, Johnstrom P, Schou M, Critchlow SE, Fawell S, Johannes JW, Leo E, Davies BR, Cosulich S, Sarkaria JN, O’Connor MJ, Hamerlik P. Preclinical Characterization of AZD9574, a Blood-Brain Barrier Penetrant Inhibitor of PARP-1. Clin Cancer Res. 2024;30(7):1338–51. doi: 10.1158/1078-0432.CCR-23-2094. PubMed PMID: 37967136.

44. Patel J, Zhou X, Gao Y, Zhao C, Li Y, Chaudhary A, Wu S, Schuster D, Liang S. <strong>A novel 18F-labeled brain penetrant PET ligand for imaging poly(ADP-ribose) polymerase-1 </strong&gt. Journal of Nuclear Medicine. 2025;66(Supplement 1):251647.

45. D’Andrea AD. Mechanisms of PARP inhibitor sensitivity and resistance. DNA Repair (Amst). 2018;71:172–6. Epub 20180823. doi: 10.1016/j.dnarep.2018.08.021. PubMed PMID: 30177437.

46. Fu X, Li P, Zhou Q, He R, Wang G, Zhu S, Bagheri A, Kupfer G, Pei H, Li J. Mechanism of PARP inhibitor resistance and potential overcoming strategies. Genes Dis. 2024;11(1):306–20. Epub 20230324. doi: 10.1016/j.gendis.2023.02.014. PubMed PMID: 37588193; PMCID: PMC10425807.

47. Noordermeer SM, van Attikum H. PARP Inhibitor Resistance: A Tug-of-War in BRCA-Mutated Cells. Trends Cell Biol. 2019;29(10):820–34. Epub 20190814. doi: 10.1016/j.tcb.2019.07.008. PubMed PMID: 31421928.

