## Supplemental FIgure 1 for "A novel ^18^F-labeled brain penetrant PET ligand for imaging poly(ADP-ribose) polymerase-1"

**Synthesis of standard compound 5**

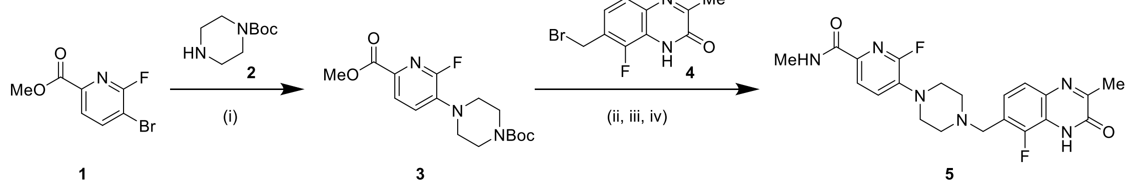


**Synthesis of brominated labeling precursor 8**

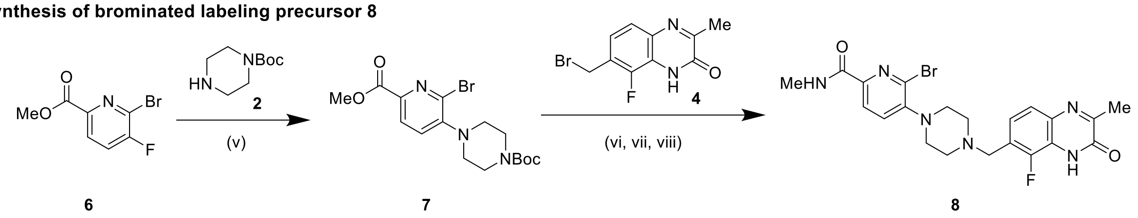


**Radiolabeling of [$^{18}$F]AZD9574**

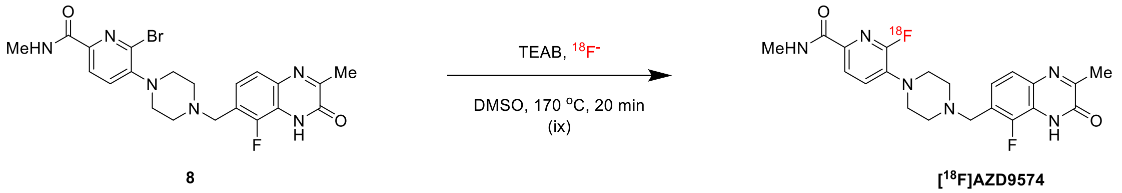


**Figure S1:** Synthesis of standard AZD9574, labeling precursor Br-AZD9574 and radiosynthesis of [$^{18}$F]AZD957 with three representative conditions and associated radiochemical yields (RCY). (i) *N*-Boc piperazine, Pd-RuG3, $Cs_2CO_3$, 1,4-dioxane, 100 ºC, 24 h, 56%; (ii) HCl in 1,4-dioxane (3 M), 6 h, rt; (iii) DIPEA, MeCN, 80 ºC ,12 h, 20%; (iv) $MeNH_2$ in THF (2 mol/L), EtOH, 80 ºC, 12 h, 55%; (v) $K_2CO_3$, DMF, 130 ºC, 48%; (vi) HCl in 1,4-dioxane (3 mol/L), 6h, rt; (vii) DIPEA, MeCN, 80ºC, 12h, 26%; (viii) $MeNH_2$ in THF (2 mol/L), EtOH, 80 ºC, 12 h, 64%. (ix) $^{18}$F anion in $^{18}$O water, TEAB (3 mg), DMSO (0.3 mL), 170 ºC 20 min.
